# The proprotein convertase BLI-4 promotes collagen secretion during assembly of the *Caenorhabditis elegans* cuticle

**DOI:** 10.1101/2023.06.06.542650

**Authors:** Susanna K. Birnbaum, Jennifer D. Cohen, Alexandra Belfi, John I. Murray, Jennifer R.G. Adams, Andrew D. Chisholm, Meera V. Sundaram

## Abstract

Some types of collagens, including transmembrane MACIT collagens and *C. elegans* cuticle collagens, are N-terminally cleaved at a dibasic site that resembles the consensus for furin or other proprotein convertases of the subtilisin/kexin (PCSK) family. Such cleavage may release transmembrane collagens from the plasma membrane and affect extracellular matrix assembly or structure. However, the functional consequences of such cleavage are unclear and evidence for the role of specific PCSKs is lacking. Here, we used endogenous collagen fusions to fluorescent proteins to visualize the secretion and assembly of the first collagen-based cuticle in *C. elegans* and then tested the role of the PCSK BLI-4 in these processes. Unexpectedly, we found that cuticle collagens SQT-3 and DPY-17 are secreted into the extraembryonic space several hours before cuticle matrix assembly. Furthermore, this early secretion depends on BLI-4/PCSK; in *bli-4* and cleavage-site mutants, SQT-3 and DPY-17 are not efficiently secreted and instead form large intracellular aggregates. Their later assembly into cuticle matrix is reduced but not entirely blocked. These data reveal a role for collagen N-terminal processing in intracellular trafficking and in the spatial and temporal restriction of matrix assembly *in vivo*. Our observations also prompt a revision of the classic model for *C. elegans* cuticle matrix assembly and the pre-cuticle-to-cuticle transition, suggesting that cuticle layer assembly proceeds via a series of regulated steps and not simply by sequential secretion and deposition.

## Introduction

Proteolytic cleavage is a common regulatory step in the assembly of extracellular matrices: Components initially enter the secretory pathway as soluble proproteins and then are cleaved in later secretory compartments or extracellularly to allow their assembly into higher order structures [1–3]. For example, mammalian fibrillar collagens undergo both N-terminal and C-terminal proteolysis in order to convert procollagen to the mature collagen that is found in extracellular fibrils [4–7]. For Type I procollagen, these cleavages depend (at least in part) on members of the ADAMTS and BMP-1/astacin proteinase families, respectively [5,8–10], and failure to appropriately cleave procollagen leads to human connective tissue disorders such as Ehlers-Danlos syndrome Type VII (ED-VII) and Osteogenesis imperfecta [4,9–12]. Some other types of collagens, including transmembrane MACIT collagens (membrane associated collagens with interrupted triple helixes) and *C. elegans* cuticle collagens, are instead N-terminally cleaved at a dibasic site that resembles the consensus for furin or other proprotein convertases of the subtilisin/kexin (PCSK) family [13–16]. However, evidence for the importance of specific PCSKs in collagen cleavage is lacking and the functional consequences of such cleavage are not well defined.

Appropriate cleavage of procollagens may facilitate formation of fibrils or other higher order matrix structures at the right place and time, in the presence of appropriate partners. In the case of transmembrane collagens, N-terminal cleavage also would release the ectodomain from the plasma membrane. Unfortunately, potential redundancy and uncertainty about the specific proteinases involved, combined with difficulties in visualizing collagen matrix *in vivo*, have made it challenging to dissect these regulatory mechanisms in most biological systems. For example, while C-terminal cleavage strongly promotes Type I collagen fibril assembly *in vitro* [5,17], there is still uncertainty about the specific role N-terminal cleavage plays in matrix assembly [3,6,7,18]. Unprocessed Type I Pro(N)-collagen can be found extracellularly in cell culture [4,19] and in morphologically abnormal fibrils in ED-VII patients [20,21], leading to the widespread view that N-terminal processing is not essential for collagen secretion or incorporation into fibrils, but rather affects specific aspects of fibril structure. It remains unclear if this is generally true for other types of collagens, including those cleaved by PCSKs.

The nematode *Caenorhabditis elegans* has an external body cuticle that consists primarily of collagens, many of which have a predicted transmembrane domain and/or an N-terminal consensus furin cleavage site (CFCS) that could be cleaved by a PCSK [13,22–24]. A new collagenous cuticle matrix is synthesized in the embryo and during each molt between larval stages, and it is always preceded by a transient “pre-cuticle” matrix that contains zona pellucida (ZP) domain proteins and other non-collagen components [25–31]. Therefore, these two matrix types must be assembled and then disassembled in the proper sequence during the molt cycle. These dynamic matrix changes are controlled in part by oscillatory gene expression programs, with pre-cuticle genes peaking relatively early in each molt cycle and different cuticle collagen genes peaking at early, intermediate, or late timepoints, consistent with an extended period of cuticle synthesis and assembly [32–34]. The molt cycle also is controlled by various post-transcriptional mechanisms such as regulated trafficking and proteolysis [35–40]. A BMP1-related astacin proteinase, DPY-31, has been implicated in C-terminal processing of SQT-3 cuticle collagen [41–43], while one or more furin/PCSKs are thought to be responsible for N-terminal processing of many cuticle collagens at the CFCS. The functional importance of N-terminal cleavage is supported by the fact that CFCS mutations in several collagens cause disruptions to cuticle structure [13,14,22].

Here we investigate the roles of *C. elegans* BLI-4, a member of the furin/PCSK family [44], in cuticle assembly. PCSK proteinases cleave secreted or transmembrane substrates immediately following dibasic sites of consensus sequence (R/K) Xn (R/K), where Xn can be 0, 2, 4, or 6 amino acids [45]. Mammals have 9 PCSK family members that differ primarily in their C-terminal domains, which are thought to confer different subcellular localization patterns and/or substrate preferences [45–47]. *C. elegans* has 4 PCSK family members, among which BLI-4 and KPC-1 appear to be the major ones expressed in external epithelial cells, while EGL-3 and AEX-5 are expressed primarily in neurons or internal epithelia [48–50]. *kpc-1, egl-3,* and *aex-5* mutants are viable and do not have any reported defects in the molt cycle or cuticle [51–53]. In contrast, *bli-4* is an essential gene, and isoform-specific *bli-4(e937)* mutants have blistered adult cuticles, making BLI-4 an excellent candidate for cleaving cuticle collagens [44]. By imaging formation of the first (L1) cuticle in developing wild-type embryos and *bli-4* or cleavage site mutants, we provide evidence that BLI-4-dependent N-terminal processing of specific cuticle collagens promotes their efficient secretion and restricts premature matrix assembly.

## Results

### Collagen secretion begins several hours before the pre-cuticle to cuticle transition

The collagenous cuticle matrix of *C. elegans* is preceded by a molecularly distinct pre-cuticle apical extracellular matrix (aECM) that it eventually replaces during each molt cycle [54]. To determine the precise timing and sequence of these events for the first pre-cuticle and cuticle, we imaged staged live embryos expressing fluorescently-tagged matrix factors (Fig. 1). Tags were located either internally (int), at the N-terminus immediately following the signal sequence (ss), or at the extreme C-terminus, as described in Table S2 and schematized in the figures below. All of these matrix fusions were generated by CRISPR/Cas9-dependent tagging of the endogenous loci and were functional based on phenotypic assays (Methods). Images were collected at the 1.5-fold stage and then at one-hour intervals thereafter, as the embryos elongated to their ultimate worm body shape.

**Figure 1.**
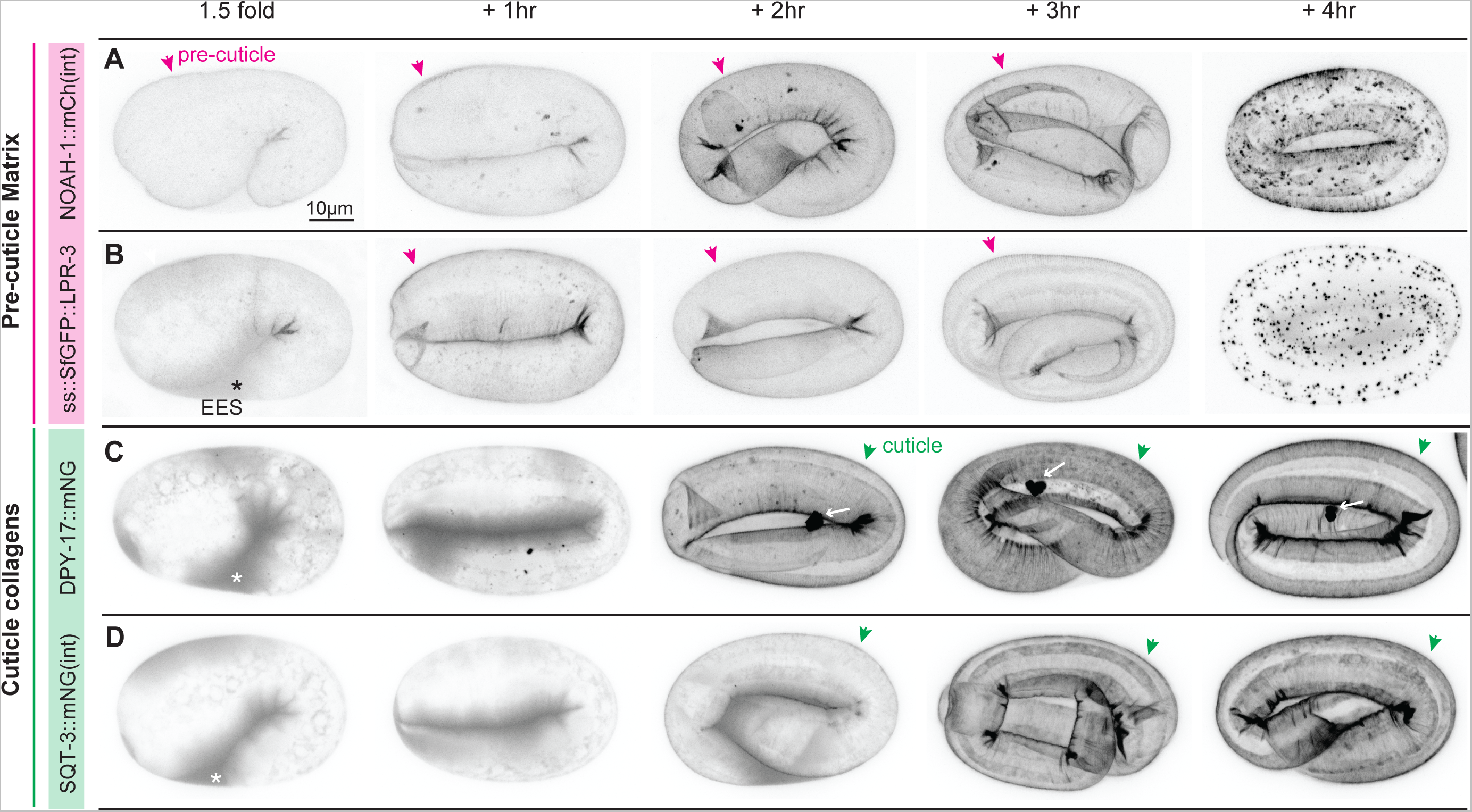
Matrix dynamics during assembly of the first pre-cuticle and cuticle. *C. elegans* embryos expressing functional pre-cuticle (A-B) or cuticle collagen (C-D) fusion proteins from endogenously-tagged loci. A) *noah-1(mc68* [NOAH-1::mCherry(int)]). B) *lpr-3(cs250* [ss::SfGFP::LPR-3]). C) *dpy-17(syb3685* [DPY-17::mNG]). D) *sqt-3(syb3691* [SQT-3::mNG(int)]). Embryos were selected at the 1.5-fold stage and then incubated for the indicated number of hours prior to imaging. Magenta arrowheads indicate pre-cuticle sheath. Green arrowheads indicate cuticle and white arrows indicate extracellular DPY-17::mNG aggregates. Asterisks indicate secreted fusion protein within the extraembryonic space (EES). All images are maximum intensity projections from confocal Z-slices, shown in inverted grayscale for clarity. Images are representative of at least 5 embryos per genotype per stage. Scale bar, 10 microns. **Related to Figure 1** **Figure S1. Pre-cuticle mCherry fusions accumulate in lysosome-like compartments following endocytosis**

The pre-cuticle or “sheath” matrix can be detected by the 1.5-fold stage and is important for proper embryo elongation beyond the 2-fold stage [25,29,55]. Consistent with that, the pre-cuticle ZP protein NOAH-1 tightly marked the pre-cuticle from before 1.5-fold to post-elongation (Fig. 1A), as previously reported [29]. The secreted lipocalin LPR-3 initially accumulated between the embryo and the eggshell, and then marked pre-cuticle beginning about an hour after NOAH-1 (Fig. 1B). Both proteins then were endocytosed and cleared. mCherry fusions but not Superfolder (Sf) GFP fusions accumulated in large lysosome-like structures (Fig. S1), suggesting that endocytosed protein eventually moved into an acidic endolysosomal compartment [56]. Together these data indicate that pre-cuticle matrix assembly occurs in a stepwise fashion. Furthermore, pre-cuticle matrix clearance involves considerable endocytosis.

Assembly of the first cuticle traditionally has been thought to begin late in embryogenesis, about 4 hours after 1.5-fold [55,57]. However, we found that tagged cuticle collagens DPY-17 and SQT-3 were secreted before the 1.5-fold stage, contemporaneously with pre-cuticle factors (Fig. 1C,D). These collagens initially accumulated between the embryo and the eggshell, began to incorporate detectably into matrix by the 2-hour timepoint (when embryos had elongated to the 3-fold stage), and appeared fully incorporated by the 3-to-4-hour timepoints. DPY-17 also consistently marked a single large extracellular aggregate that appeared in the extracellular space concomitant with matrix incorporation, consistent with some change in its molecular properties at this time (Fig. 1C). In summary, the transition from pre-cuticle to cuticle begins during embryo elongation and components of both types of matrices transiently coexist, with the pre-cuticle disassembling once embryo elongation is complete. Furthermore, collagen secretion occurs earlier than previously thought, yet detectable collagen matrix incorporation occurs 2-3 hours afterwards, consistent with post-transcriptional (and possibly post-secretory) controls of cuticle matrix assembly.

### BLI-4 is widely expressed in external epithelia and localizes to intracellular compartments and the extraembryonic space

The PCSK BLI-4 is a strong candidate for cleaving cuticle collagens to promote matrix assembly [22,44]. Existing single cell RNA sequencing (scRNAseq) data from embryos [49] revealed *bli-4* expression in external epithelial cells (e.g. those lined by pre-cuticle/cuticle), including the epidermis and various interfacial tubes and glia, as well as in the foregut (pharynx), intestine and germline (Fig. S2). Consistent with this, a *bli-4* transcriptional reporter [58] also showed widespread epithelial expression, including in both the lateral (seam) and major (hyp7) epidermis and in the excretory duct and pore tubes (Fig. 2A). A functional BLI-4::SfGFP(int) fusion protein, tagged at the protease domain and expressed from the endogenous locus, marked sub-apical intracellular compartments of external epithelia at the 1.5-fold stage and beyond (Fig. 2B). BLI-4::SfGFP(int) also was faintly visible in the extraembryonic space (Fig. 2B) and it appeared transiently within the lumen of the foregut at the 1.5+4 hr timepoint (Fig. 2C). In summary, BLI-4 is expressed in external epithelia before and during cuticle assembly, and it appears both intracellular and extracellular.

**Figure 2.**
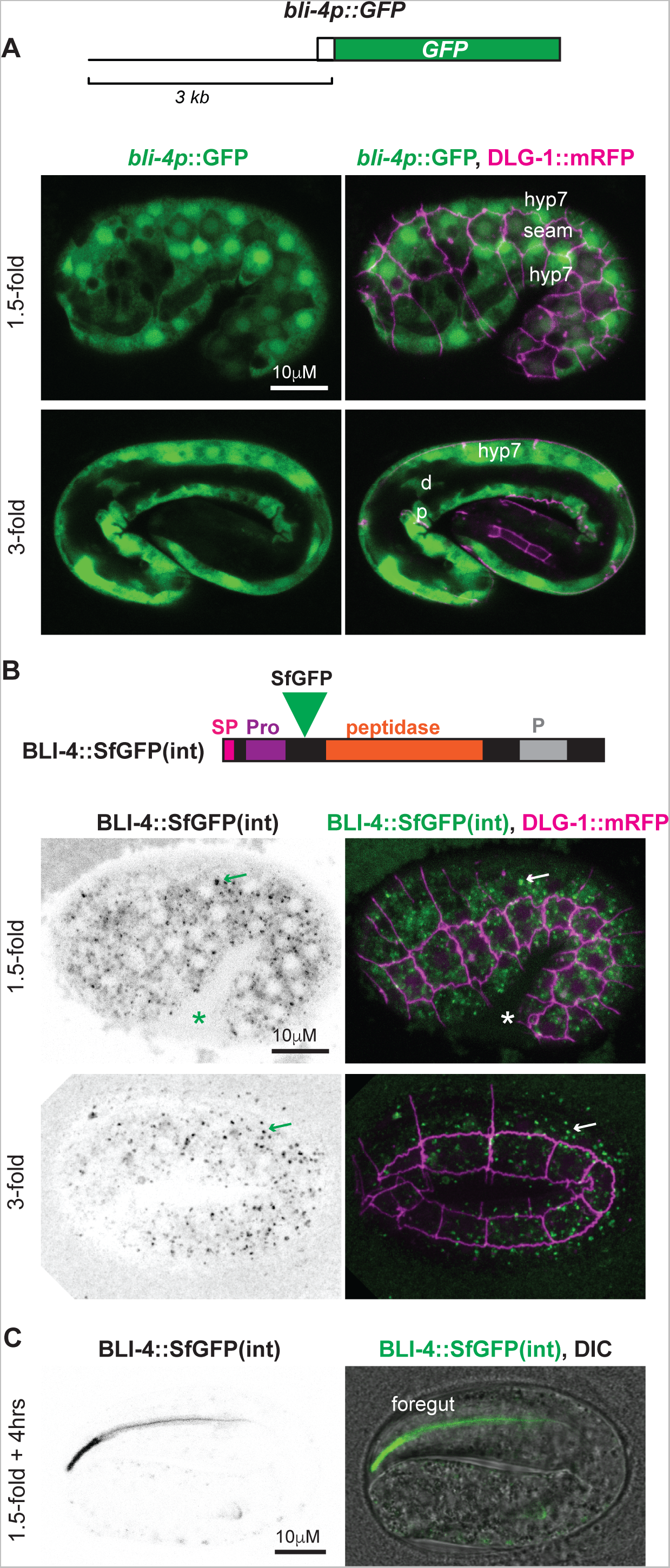
BLI-4 localizes to intracellular and extracellular compartments. A,B,C) Transcriptional and translational reporters reveal *bli-4* expression throughout pre-cuticle and cuticle assembly. Animals also express the epithelial junction marker DLG-1::RFP (*mcIs46*, magenta) to aid in cell identification and Z-depth assessment. Images are maximum intensity projections from confocal Z-stacks and representative of at least n=5 animals examined. A) A *bli-4pro::GFP* transcriptional reporter (*sEx11763*, green) [58] is broadly expressed in external epithelia, including in hyp7, seam cells, and in the excretory duct (d) and pore (p). B) An endogenous BLI-4::SfGFP(int) translational fusion (*syb5321*) marks intracellular puncta within epithelia and is faintly detectable within the extraembryonic space (EES). Single channel images are shown in inverted grayscale for clarity. Asterisk, fusion protein detected in the extraembryonic space. Arrow, intracellular puncta. CRISPR/Cas9 was used to insert SfGFP between the BLI-4 prodomain (Pro) and peptidase domain, as indicated (Table S2). The schematic shows the short isoform BLI-4f (Genbank NP_001360008.1), but all isoforms should be tagged. ss, signal sequence. P, P domain. C) BLI-4::SfGFP(int) transiently accumulated in the foregut at the 1.5 + 4hr timepoint.

### Generation of *bli-4* null and isoform-specific mutants

The *bli-4* gene has many splice isoforms that differ in their 3’ exons; the resulting proteins all share the N-terminal peptidase domain but differ in the presence or absence of other domains (Fig. 3A,B) [44]. These isoforms may have different substrate specificities. For example, *bli-4(e937)* (hereafter *bli-4(ΔBLI))* is a deletion removing exons unique to isoforms a, e, g, and h; these mutants are viable but have a blistered (Bli) adult cuticle, suggesting failure to process key substrates unique to that stage [44]. BLI-4 isoforms c and d contain a cysteine-rich domain (CRD) similar to that found in mammalian furin, PCSK5, and PCSK6 (Fig. 3A). The PCSK5 CRD can confer cell surface anchoring via binding to heparan sulfate proteoglycans [46,47], and was therefore proposed to affect substrate specificity. Based on an analysis of 3’ end reads from embryo scRNAseq data [49], the *bli-4a, d* and *f/g* isoforms are detectably expressed in the embryo. *bli-4d* is by far the most highly expressed isoform in the embryonic epidermis and is also detected at lower levels in other tissues (Fig. S2). Isoform *bli-4a* is expressed primarily in the germline and isoforms f/g are expressed most strongly in pharyngeal epithelial cells, though both are also detected in epidermis to a lesser degree (Fig. S2). The predicted sizes of these BLI-4 isoforms are consistent with the most prominent bands observed on Western blots of lysates from BLI-4::SfGFP(int)-expressing embryos (Fig. 3C,D).

**Figure 3.**
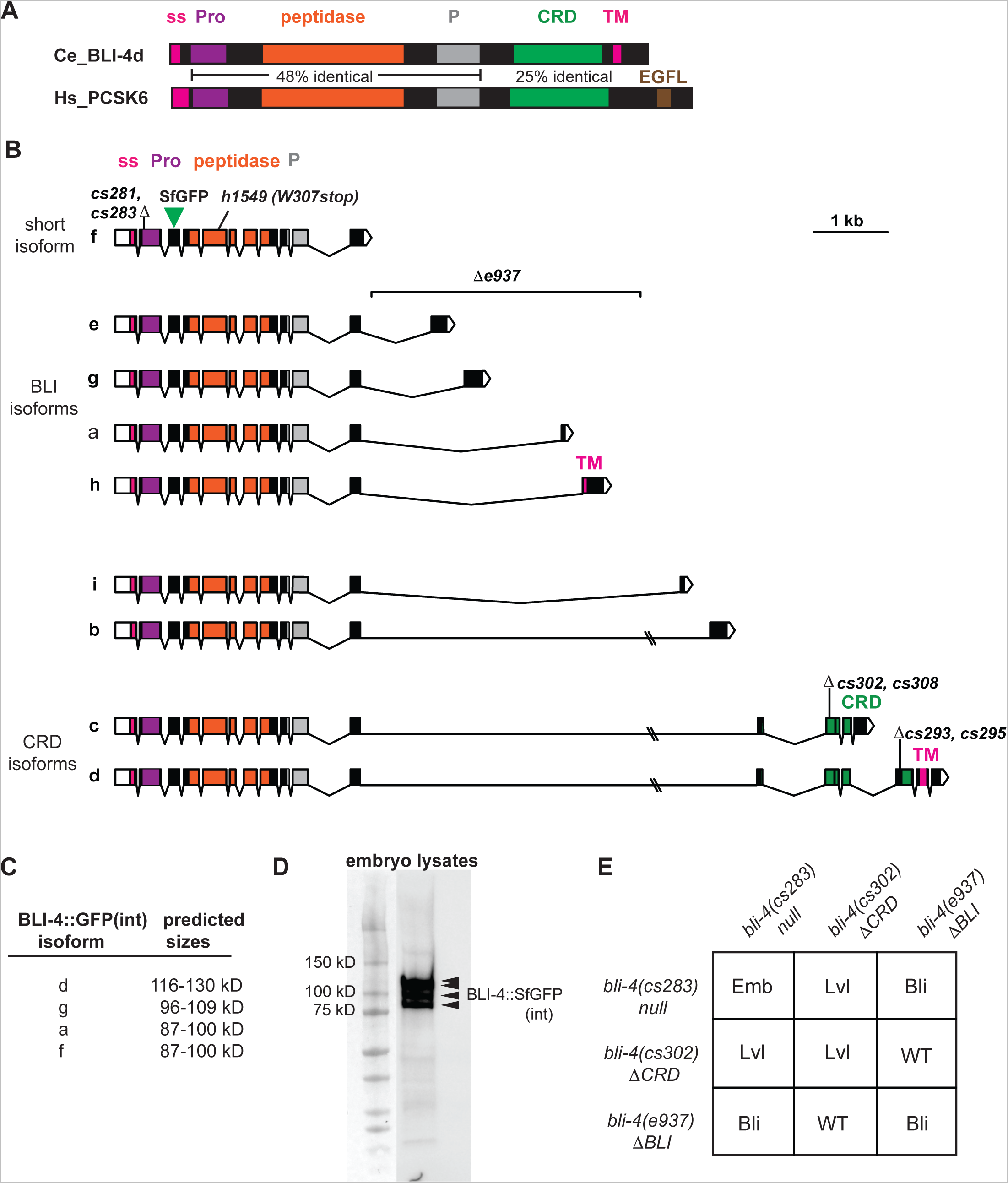
Generation of *bli-4* null and isoform-specific alleles. A) *C. elegans* BLI-4d (Genbank NP_001021543.1) protein schematic and comparison to human PCSK6 (Genbank BAA21625.1). The % identity at the amino acid level is listed between relevant domains. Like other PCSK family members, both proteins have a signal peptide (SP) and prodomain (Pro) that are removed during trafficking, followed by the peptidase domain and an associated P domain thought to assist with its folding and stability [45]. BLI-4d and PCSK6 also share a cysteine-rich domain (CRD). BLI-4d also has a transmembrane (TM) domain but it lacks the EGF-like (EGFL) domain found in PCSK6. B) *bli-4* gene isoforms and mutant alleles. Colors indicate encoded protein domains, as in A. Isoforms are arranged by mutant groups. *cs281* and *cs283* are 1-2nt deletion/frameshift mutations in exon 2, which is shared among all *bli-4* isoforms. *e937* is a 3,325 bp deletion that removes intronic sequences and exons associated with isoforms a, e, g, and h [44]. *cs302* and *cs308* are 4-19 nt indel/frameshift mutations in the first exon unique to isoforms c and d. *cs293* and *cs295* are identical 5nt deletion/frameshift mutations in the first exon unique to isoform d. See Table S2 for specific allele sequences. C) Predicted sizes of major embryonically-expressed BLI-4::SfGFP(int) fusion proteins before and after removal of the Pro domain. Sizes were estimated based on the isoform sequence using https://www.bioinformatics.org/sms/prot_mw.html. See also Figure S2 for data regarding isoform expression in the embryo. D) Western blot of lysates from BLI-4::SfGFP(int) expressing embryos. Arrowheads indicate four major bands between 80 and 130 kD. Blot is representative of 3 replicates. E) Summary of complementation test results. *bli-4(cs281)* failed to complement both *bli-4(cs302)* (n=59) and *bli-4(e937)* (n=120) for the larval lethal (Lvl) and adult Blister (Bli) phenotypes, respectively, while *bli-4(cs302)* complemented *bli-4(e937)* (n=186). Balancer *hT2 [bli-4(e937) let-?(q782) qIs48] (I;III)* was used as the *bli-4(e937)-*containing chromosome. **Related to Figure 3:** **Figure S2. *bli-4* isoform expression in the embryo**

To address the overall roles of BLI-4, and roles of the two CRD-containing isoforms specifically, we used CRISPR/Cas9 to generate new *bli-4* null and isoform-specific alleles in an isogenic strain background (Fig. 3B, Table S2, Methods). Two null alleles (*cs281* and *cs283,* hereafter *bli-4(-)*) were made by targeting exon 2, upstream of the peptidase domain; these cause frameshifts that should remove all splice isoforms. Two isoform c- & d-specific mutants (*cs302* and *cs308,* hereafter *bli-4(ΔCRD)*) and two isoform d-specific mutants (*cs293* and *cs295*) were made by targeting exons unique to those isoforms. All mutants were recessive lethal but could be rescued with a fosmid-based transgene containing the entire *bli-4* genomic locus (Figs. 3E, 4, 5). *bli-4(-)* failed to complement the isoform-specific alleles, as expected, whereas *bli-4(ΔCRD)* and *bli-4(ΔBLI)* complemented each other (Figs. 3E, 4B).

**Figure 4.**
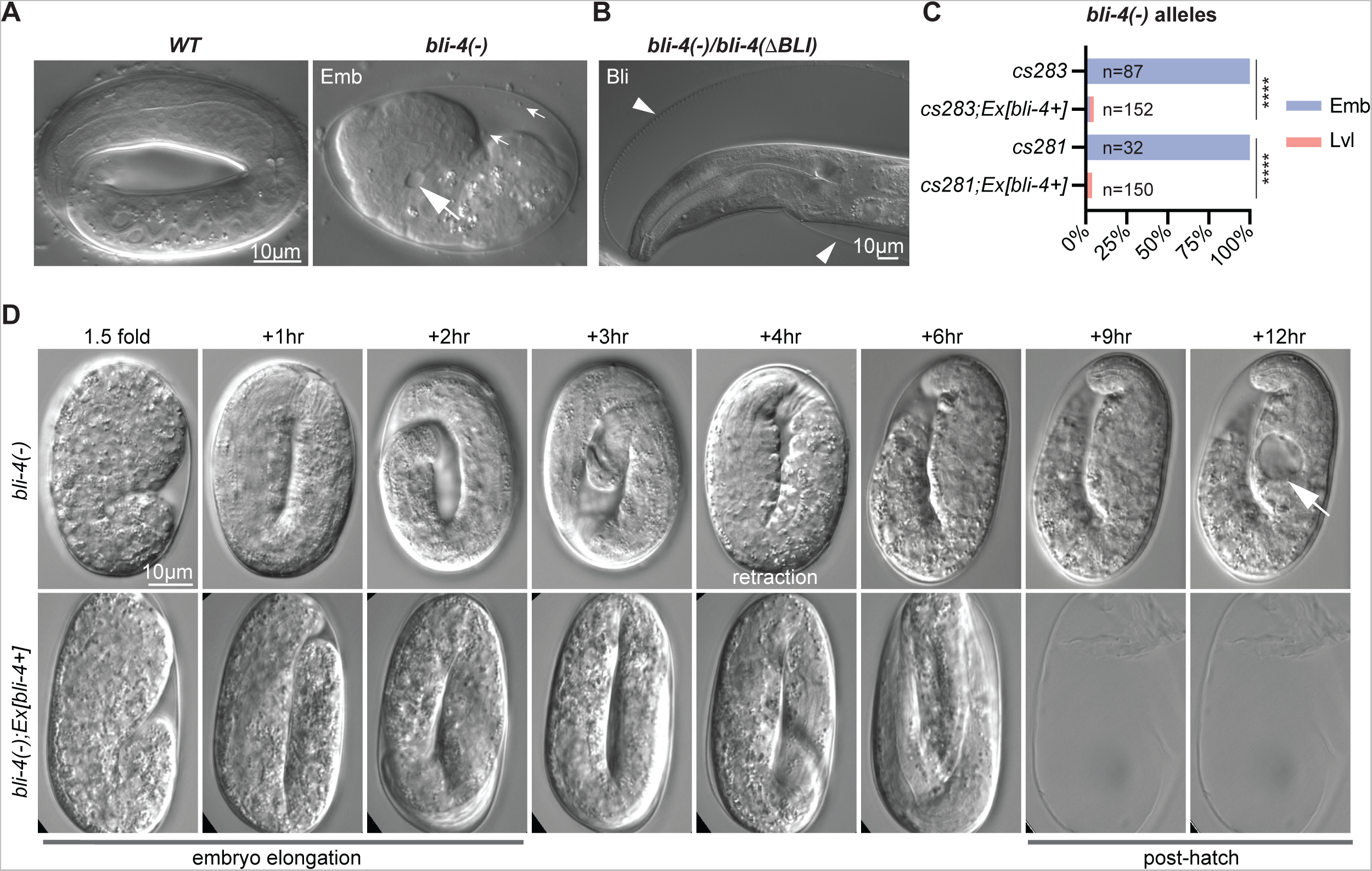
*bli-4* null mutants retract and collapse after embryonic elongation. A) In contrast to a wildtype 3-fold embryo (left), *bli-4*(-) embryos arrest as misshapen masses with excretory tube dilations (large arrows) and extracellular debris (small arrows). B) *bli-4(-)* allele *cs283* failed to complement *bli-4(e937)* for the adult Blister (Bli) phenotype (n=120). Balancer *hT2 [bli-4(e937) let-?(q782) qIs48] (I;III)* was used as the *bli-4(e937)-*containing chromosome. Arrowheads point to blistered cuticle. C) Phenotype quantitation and rescue of *bli-4(-)* mutants. *Ex(bli-4+)* corresponds to transgene *csEx919,* which contains fosmid WRM069bE05. ***P<0.0001, Fisher’s Exact Test. D) Stills from time-lapse imaging of *bli-4(cs281)* and rescued siblings at 12°C. *bli-4(-)* embryos initially elongated but then began retracting after the 3-hour timepoint (220-330 minutes, n=5). Arrow indicates excretory tube dilation. **Related to Figure 4:** **Supplementary videos 1 and 2**

**Figure 5.**
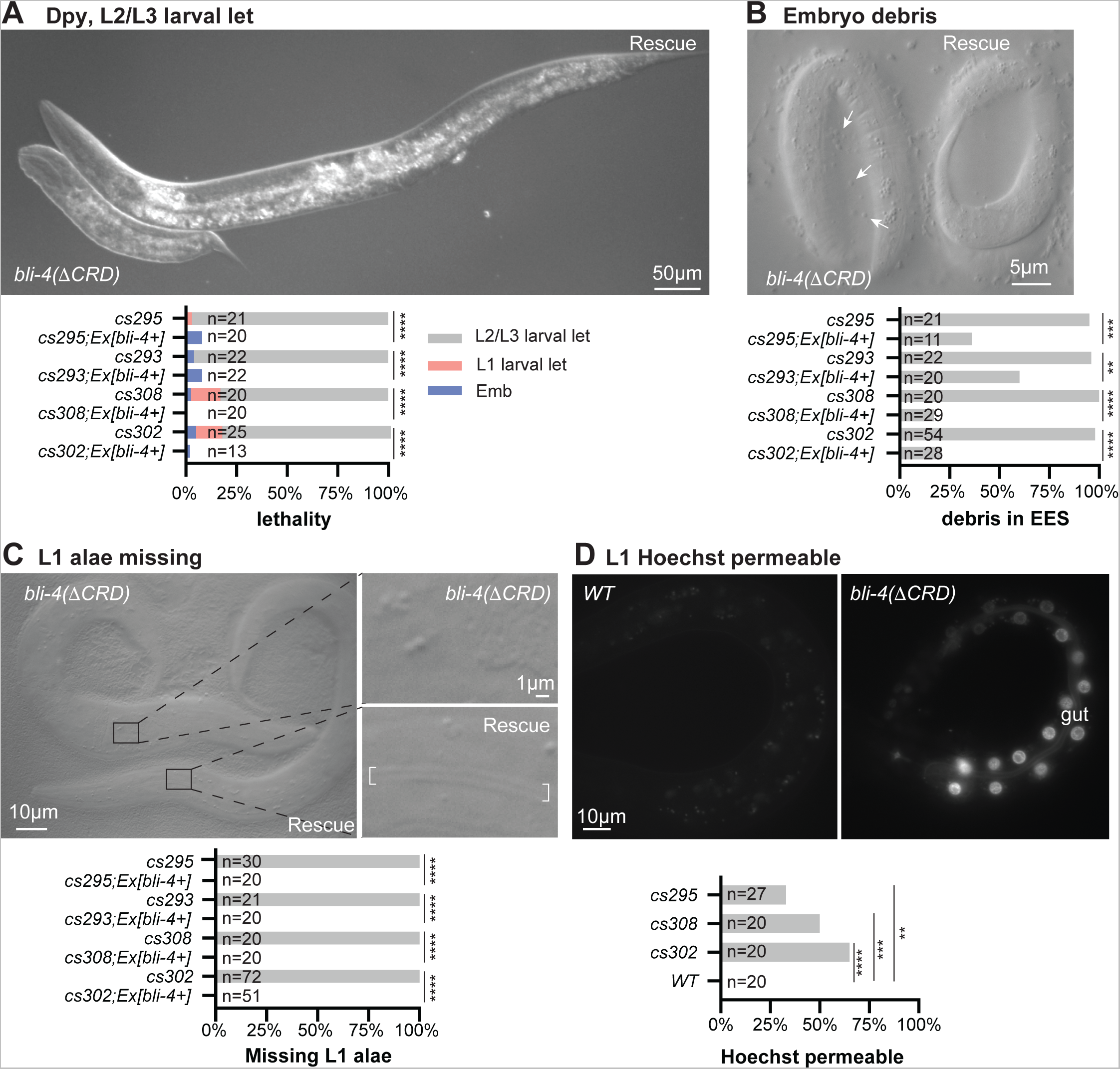
*bli-4* CRD isoform mutants arrest as Dumpy larvae with cuticle defects. A-D) *bli-4(ΔCRD)* isoform mutant phenotypes. Only images of *bli-4(cs302)* and rescued siblings are shown. Phenotype quantitation and rescue data for all alleles are shown below. The *bli-4+* rescue transgene is *csEx919*. *P<0.01, **P<0.001, ***P<0.0001, Fisher’s Exact test. A) Most mutants arrest as Dumpy L2 or L3 larvae. Larvae in image are 48hr after egg lay (AEL). B) Embryos accumulate extracellular debris (small arrows) between the embryo and the eggshell. Embryos are 1.5 fold + 3-4 hours old. C) L1 larvae are slightly Dumpy and entirely lack cuticle alae ridges. Bracket indicates position of alae in the rescued sibling. D) Whereas *wild-type* L1 larvae have a permeability barrier that excludes Hoechst dye (left), many *bli-4(ΔCRD)* mutants had strong staining in the gut epithelium. Staining was not apparent in the epidermis, suggesting a gut-specific barrier defect.

### *bli-4* null mutants arrest as retracted embryos following elongation

*bli-4(-)* mutants were embryonic lethal (Emb), as previously reported for other null alleles (Fig. 4A,C) [44]. Mutant embryos arrested as disorganized masses with occasional excretory tube dilations and evidence of debris between the embryo and the eggshell (Fig. 4A). Although the arrested embryos appeared unelongated, timelapse imaging revealed that mutant embryos did elongate initially, but then retracted and collapsed soon afterwards, about 4 hours after the 1.5-fold stage (Fig. 4D). Notably, retraction occurred near the time when the cuticle replaces the pre-cuticle (Fig. 1) and specifically resembled that previously reported for mutants lacking the early cuticle collagen SQT-3 [55].

### *bli-4(****Δ****CRD)* mutants arrest as Dumpy larvae with abnormal cuticles

The four *bli-4(ΔCRD)* or *bli-4(d)* mutants all appeared less severe than the null but similar to each other, with variable larval arrest (Lvl) at the L1, L2, or L3 stage (Fig. 5A). Despite the accumulation of debris during embryogenesis (Fig. 5B), the majority of these mutants elongated and hatched (Fig. 5A). Mutants appeared only mildly Dumpy (Dpy) as L1 larvae, but they completely lacked the alae ridges typical of the L1 cuticle (Fig. 5C). The L1 cuticle retained its barrier function to exclude Hoechst dye, but the gut permeability barrier appeared defective (Fig. 5D). *bli-4(ΔCRD)* mutants became more severely Dpy by the time of L2 or L3 arrest (Fig. 5A), suggesting that the CRD isoforms are particularly important for processing substrates during the early larval stages. This strong Dpy phenotype resembles that of many cuticle collagen mutants [23,59,60].

### BLI-4 promotes secretion and cuticle incorporation of collagens SQT-3 and DPY-17

Next, we examined our matrix fusions in *bli-4* mutant backgrounds. Although many ZP proteins are cleaved at a C-terminal CFCS site before matrix incorporation [2], we did not note any obvious change in the pre-cuticle appearance of a NOAH-1 fusion between wild type and *bli-4* mutants (Fig. S3). Therefore, protein trafficking was not generally disrupted. However, we did find significant differences in the appearance of both cuticle collagens (Figs. 6,7).

**Figure 6.**
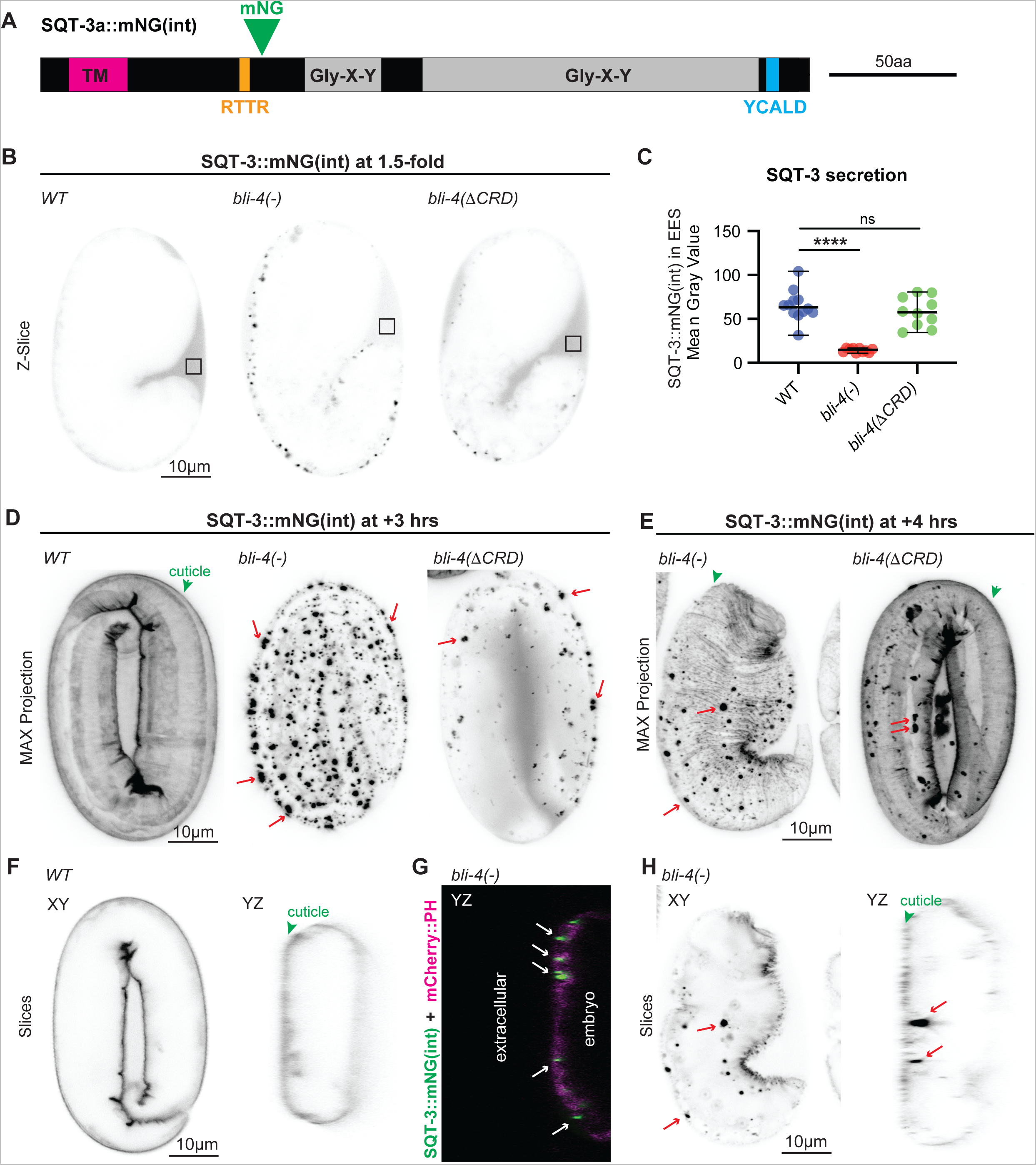
BLI-4 promotes secretion of SQT-3 cuticle collagen. A) Schematic diagram of the SQT-3a::mNG(int) fusion, showing Gly-X-Y collagen domains (gray) and N-terminal CFCS (RTTR, orange). SQT-3 lacks a signal peptide, but isoform A (shown) contains a predicted transmembrane (TM) domain (magenta). Another isoform, SQT-3b, begins after this domain and should also be tagged. A predicted DPY-31-dependent cleavage site (blue) near the C-terminus is also shown. B) SQT-3::mNG(int) is poorly secreted in *bli-4* null mutants. Single confocal Z-slices of 1.5-fold embryos expressing SQT-3::mNG. Box indicates extra-embryonic region analyzed in C. Images shown are representative of at least 10 images per genotype. C) Quantitation of SQT-3::mNG(int) signal in the extraembryonic space (EES) of *WT* vs. *bli-4* 1.5-fold embryos. Fluorescence intensity was measured within 2×2 micron boxes of single confocal Z-slices, as in B. Each circle represents data from a single animal. ***, P<0.0001, Mann-Whitney U test. D-E) SQT-3::mNG(int) forms aggregates (arrows) in *bli-4* mutants. Maximum intensity projections from confocal Z-stacks of embryos at the 1.5-fold + 3 hour (D) or 4-hour (E) stage. By the latter stage *bli-4* null mutants collapsed with minimal SQT-3::mNG(int) matrix incorporation, whereas *bli-4(cs302)* mutants showed significant incorporation into the cuticle (green arrowhead). Images shown are representative of at least 10 embryos per genotype. F-H) Orthogonal views of *WT* (F) and *bli-4(cs281)* (G, H) embryos expressing SQT-3::mNG(int). Embryo in (G) also contains a membrane marker *let-653pro::*mCherry::PH (*csIs98*). Aggregates (arrows) appear to be intracellular or to span the plasma membrane. WT embryo in (F) is the same as that shown in (D); *bli-4* embryo in (H) is the same as that shown in (E).

**Figure 7.**
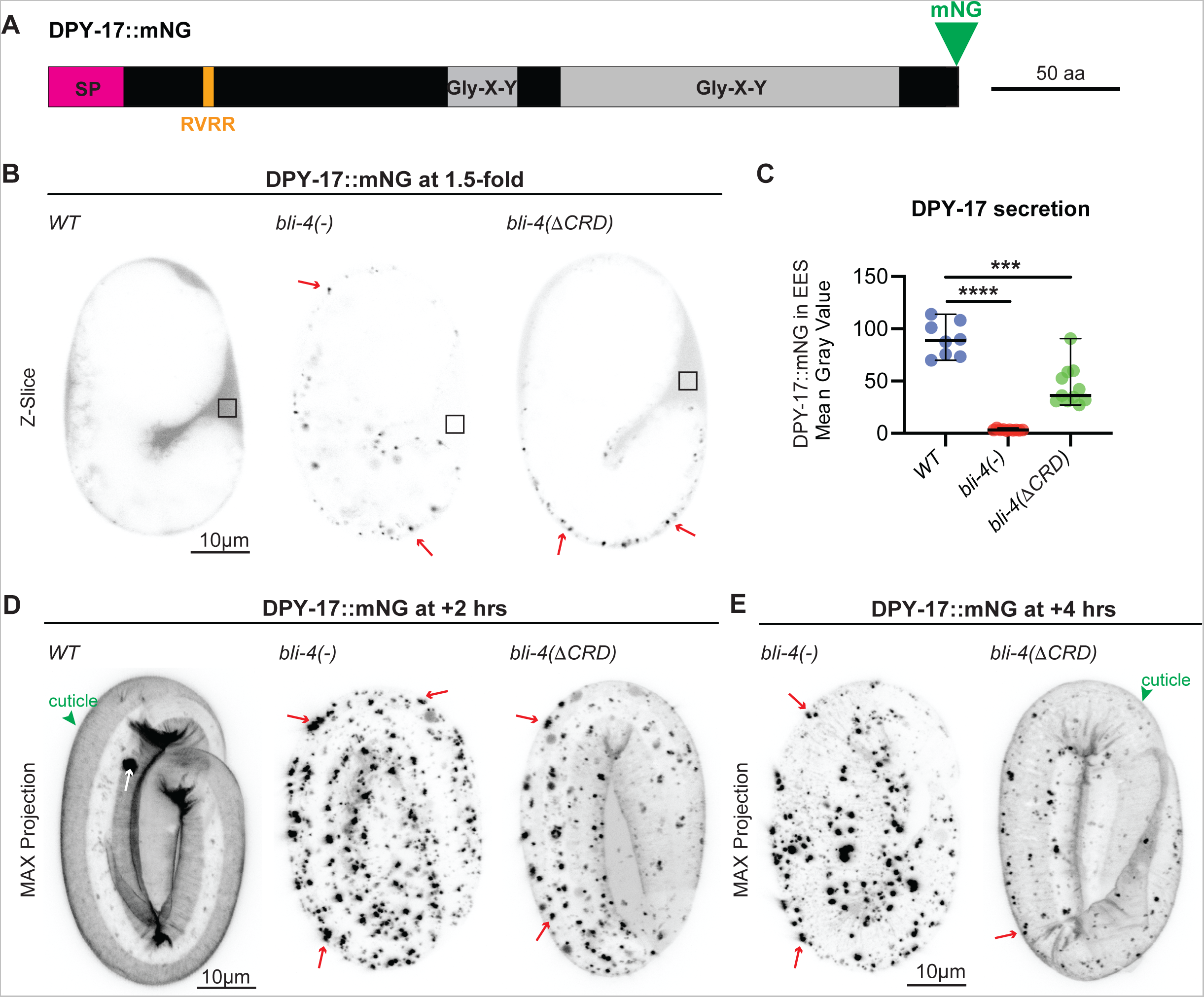
BLI-4 promotes secretion of DPY-17 cuticle collagen. A) Schematic diagram of the DPY-17::mNG fusion, showing Gly-X-Y collagen domains (gray), signal peptide (SP, magenta) and N-terminal CFCS (RVRR, orange). B) DPY-17::mNG is poorly secreted in *bli-4* null mutants. Single confocal Z-slices of 1.5-fold embryos expressing DPY-17::mNG. Box indicates extra-embryonic region analyzed in C. Images shown are representative of at least 10 images per genotype. C) Quantitation of DPY-17::mNG signal in the extraembryonic space (EES) of *WT* vs. *bli-4* 1.5-fold embryos. Fluorescence intensity was measured within 2×2 micron boxes of single confocal Z-slices, as in B. Each circle represents data from a single animal. ***, P<0.0001, Mann-Whitney U test. D,E) DPY-17::mNG forms aggregates (arrows) in *bli-4* mutants. Maximum intensity projections from confocal Z-stacks of embryos at the 1.5-fold + 2 hour (D) and 4 hour (E) stages. By the latter stage *bli-4* null mutants collapsed with minimal DPY-17::mNG matrix incorporation, whereas *bli-4(cs302)* mutants showed significant incorporation into the cuticle (green arrowhead). Images shown are representative of at least 10 embryos per genotype.

SQT-3a is a predicted transmembrane collagen with a CFCS, although a shorter SQT-3b isoform begins after the transmembrane domain and could be secreted (Fig. 6A). DPY-17 is a secreted collagen with an N-terminal CFCS (Fig. 7A). Quite dramatically, in *bli-4(-)* mutants, both SQT-3::mNG(int) and DPY-17::mNG were poorly secreted and failed to accumulate in the extra-embryonic space or to incorporate efficiently into the cuticle (Figs. 6B,C; 7B,C). Instead, both collagens formed large aggregates at or near the apical plasma membrane, with most aggregates appearing at least partly intracellular when compared to the cuticle surface or an mCherry::PH membrane marker (Figs. 6D-H, 7D,E). Aggregates were visible by the 1.5-fold stage (Figs. 6B, 7B), several hours before normal matrix incorporation (Fig. 1C,D). These data indicate that BLI-4 promotes initial solubility and secretion of these two collagens.

*bli-4(ΔCRD)* mutants had less severe defects in collagen secretion; SQT-3 and DPY-17 fusions still formed some intracellular aggregates, but they were at least partly secreted and eventually incorporated into the cuticle, albeit with delayed timing (Fig. 6C-E; 7C,D). SQT-3 is the only known cuticle collagen required for *C. elegans* embryo viability, being critical to maintain embryo shape after elongation [24,55]; its partial secretion and incorporation in *bli-4(ΔCRD)* mutants but not *bli-4(-)* mutants can explain the different arrest points of these mutants. These data are consistent with a role for multiple BLI-4 isoforms in cleavage of these collagens.

### SQT-3 and DPY-17 are mutually dependent on each other for secretion

Although *dpy-17* null mutants have a less severe Dpy phenotype than *sqt-3* null mutants, prior data suggested that DPY-17 and SQT-3 function together and that DPY-17 is required for efficient SQT-3 secretion [42,61]. We were able to confirm this result; in *dpy-17(-)* mutants, SQT-3 accumulated cytosolically and in a halo pattern surrounding epidermal nuclei (Fig. 8A,B), suggesting retention in the endoplasmic reticulum. Nevertheless, a small amount of SQT-3 did eventually incorporate into the cuticle (Fig. 8C), potentially explaining why *dpy-17* mutants are able to survive. Conversely, we found that SQT-3 is required for both DPY-17 secretion and overall protein accumulation; DPY-17 was barely detectable in *sqt-3(-)* mutants and all visible protein was intracellular (Fig. 8D,E). These data support the model that DPY-17 and SQT-3 travel together through the secretory pathway and suggest that *bli-4* loss could affect DPY-17 and SQT-3 both directly and indirectly via effects on the other (Fig. 8F).

**Figure 8.**
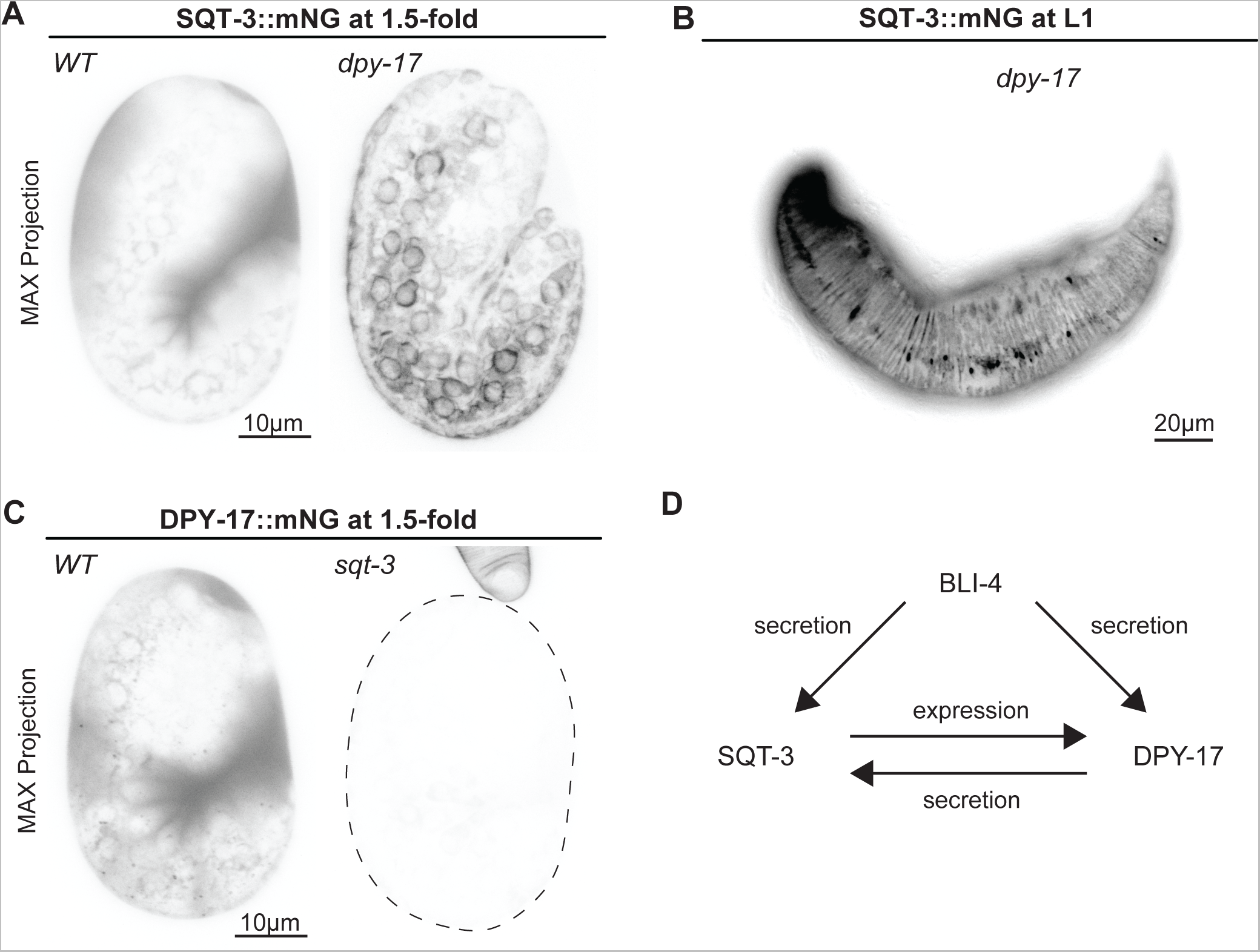
SQT-3 and DPY-17 collagens are mutually dependent for secretion. A, B) Maximum intensity projections of embryos (A) or L1 larvae (B) expressing SQT-3::mNG. A) *dpy-17(e164)* mutant embryos retained most SQT-3::mNG intracellularly; B) L1s showed some residual incorporation. Images shown are representative of at least 5 embryos per genotype. C) Maximum intensity projections of embryos expressing DPY-17::mNG at the 1.5-fold stage. DPY-17::mNG was barely detectable in *sqt-3(e2924ts)* mutants, even at the permissive temperature of 15° C. D) Model summarizing SQT-3 and DPY-17 relationship and possible direct and indirect effects of BLI-4 on each.

### CFCS mutations in SQT-3 and DPY-17 mimic loss of *bli-4*

The dramatic aECM defects described above are consistent with roles for BLI-4 in the N-terminal processing of multiple cuticle collagens. Unfortunately, because *bli-4* mutants are lethal, it is difficult to collect large numbers of mutant embryos and we have not found appropriate *bli-4* knockdown and Western blot conditions to directly test if BLI-4 is required for collagen cleavage. Instead, we used CRISPR/Cas9 to mutate the predicted BLI-4-dependent cleavage sites from RxxR to AxxA within the endogenous SQT-3::mNG(int) and DPY-17::mNG fusions (see Figs. 6,7). These mutations mimicked loss of *bli-4*, causing aggregation and intracellular retention of the mutant proteins (Fig. 9A,B). Some mutant collagen eventually incorporated into the L1 cuticle, but matrix structures appeared abnormal and larvae exhibited a severe Dpy phenotype (Fig. 9C,D). These data strongly support the model that BLI-4 directly cleaves both collagens to promote their secretion.

**Figure 9:**
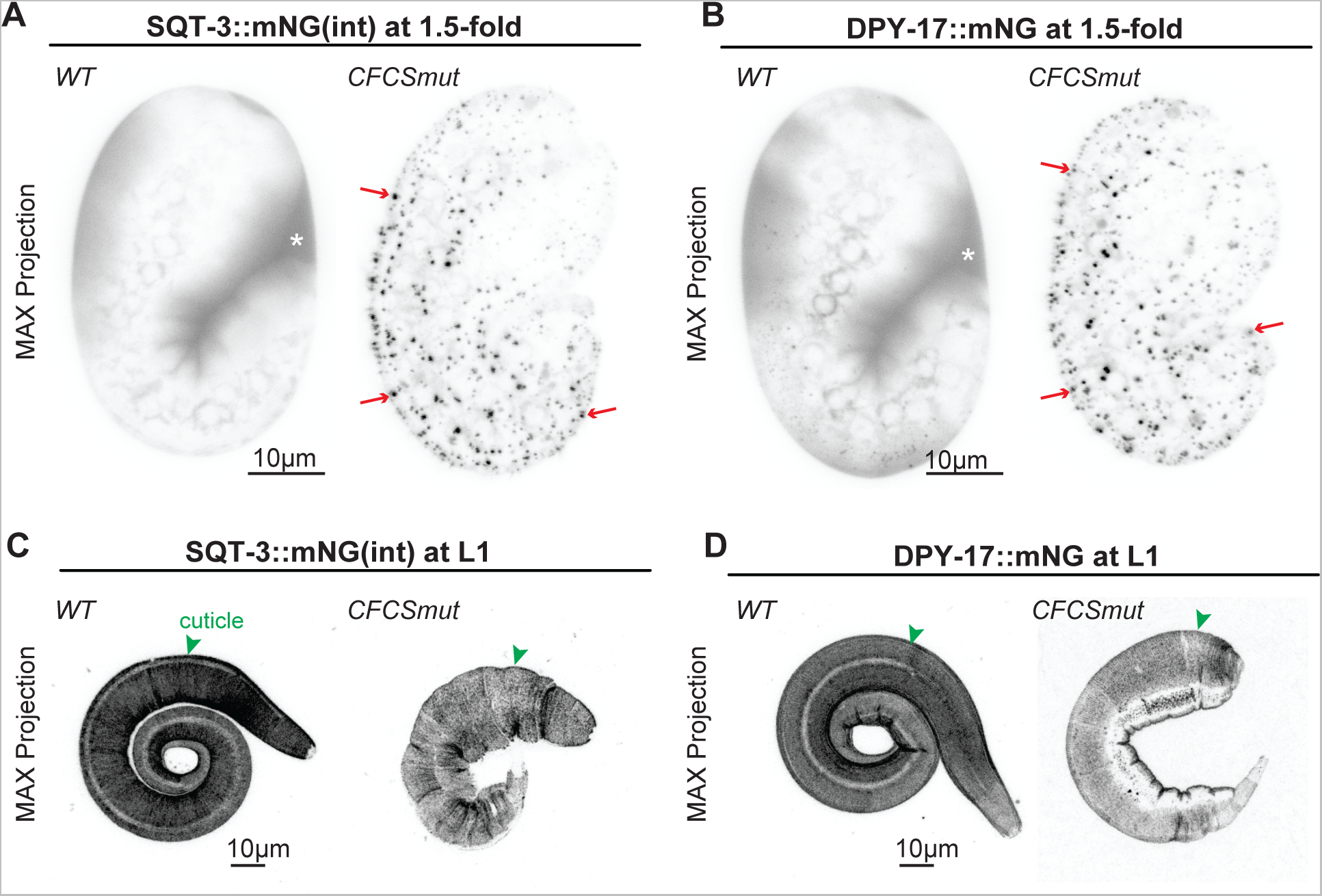
CFCS mutants for SQT-3 and DPY-17 form intracellular aggregates. Maximum intensity projections of 1.5-fold embryos (A,B) or L1 larvae (C,D) expressing A,C) SQT-3::mNG(int) vs. SQT-3(R79A,R82A)::mNG(int)) and (B,D) DPY-17::mNG vs. DPY-17(R61A,R63A,R64A)::mNG. Although poorly secreted, the CFCS mutant collagens did eventually incorporate into cuticle. Images shown are representative of at least 10 embryos and 5 L1s per genotype.

### SQT-3 CFCS mutant aggregation occurs independently of DPY-31 astacin

DPY-31, a BMP1-related astacin protease, previously was proposed to cleave the SQT-3 C-terminus to promote matrix assembly [41–43]. We confirmed that, in *dpy-31* mutants, SQT-3 was still efficiently secreted and did not form aggregates, and its cuticle incorporation was delayed compared to wild type (Fig. 10A). However, SQT-3 eventually incorporated into the cuticle of *dpy-31* mutants (Fig. 10B,D), suggesting either that C-terminal cleavage is not absolutely required or that other proteases can also execute this cleavage (Fig. 10E). Furthermore, *dpy-31* loss did not suppress the aggregation of SQT-3 CFCS mutants (Fig. 10C,D); therefore, aggregation is not due to premature cleavage by DPY-31. Instead, retention of the SQT-3 N-terminus may allow inappropriate interactions with other unknown factors along the secretory pathway, leading to intracellular aggregation (Fig. 10E).

**Figure 10:**
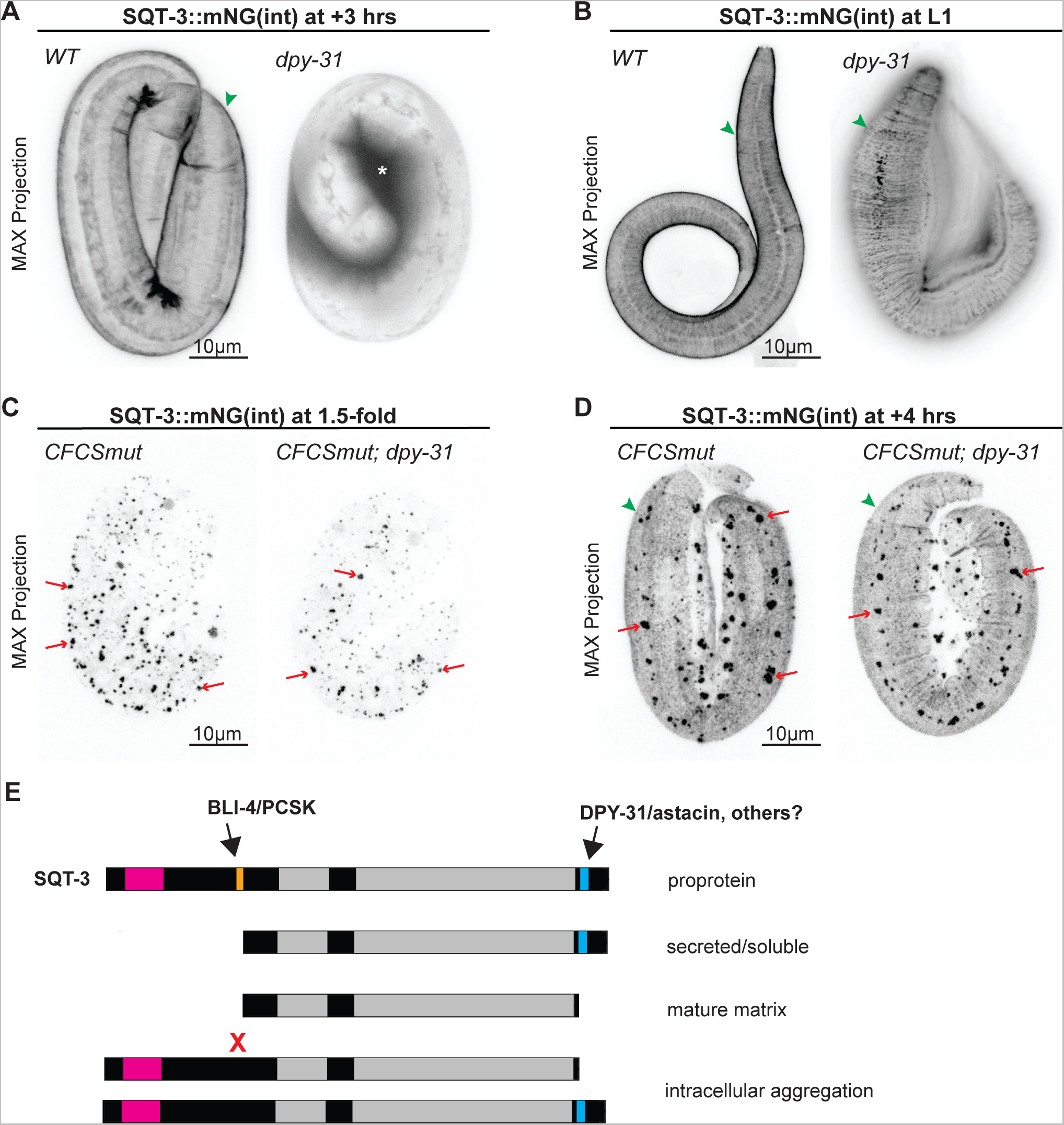
SQT-3 CFCS mutant aggregates form independently of DPY-31/astacin. Maximum intensity projections of WT vs. *dpy-31(e2770)* mutants expressing A,B) SQT-3::mNG(int) or C,D) CFCS mutant SQT-3(R79A,R82A)::mNG(int)). DPY-31 promotes timely SQT-3 matrix incorporation (A) and proper cuticle structure (B), but its removal does not suppress the SQT-3 CFCS aggregation defect (C,D). Images shown are representative of at least 5 animals per genotype. E) Model for sequential cleavage of SQT-3 collagen by BLI-4/PCSK and DPY-31/astacin. BLI-4 promotes secretion of soluble forms of SQT-3, while DPY-31 and possibly other astacin proteases promote mature matrix assembly. Preventing BLI-4-dependent N-terminal cleavage (X) leads to intracellular aggregation. This aggregation occurs independently of DPY-31, but could reflect premature matrix assembly due to inappropriate modification by other assembly-promoting factors such as other astacin proteases, proline hydroxylases [81], or tyrosine cross-linking enzymes [67,68].

## Discussion

N-terminal processing is an important step in the maturation of most collagens and is generally thought to affect fibril structure. Here we provide evidence for an alternative role of N-terminal processing in *C. elegans* cuticle collagens: to allow secretion of soluble forms of collagen and prevent premature matrix assembly and aggregation (Fig. 10E). Using new collagen fusion knock-ins that allowed us to visualize *C. elegans* cuticle assembly in live embryos, we showed that collagen secretion into the extraembryonic space precedes cuticle assembly by several hours. Loss of *bli-4* PCSK prevents efficient secretion of two early cuticle collagens, SQT-3 and DPY-17, causing them to aggregate within apical compartments of epidermal cells and reducing (but not completely blocking) their later assembly into cuticle matrix. Mutation of the predicted BLI-4-dependent cleavage sites causes similar aggregation defects. These data demonstrate a role for collagen N-terminal processing in intracellular trafficking and in the spatial and temporal restriction of matrix assembly *in vivo*.

### Temporal control of cuticle assembly and the pre-cuticle to cuticle transition

The *C. elegans* pre-cuticle and cuticle are molecularly distinct matrices that coat external epithelia at different stages of development. The pre-cuticle is present at earlier stages and is required for initial embryo elongation, while the cuticle eventually replaces it and is responsible for maintaining embryo shape [25,29,55]. Consistent with those defined roles, we showed here that the transition to cuticle matrix happens during and shortly following embryo elongation. Surprisingly, prior to this transition, at least some cuticle collagens are present in the extra-embryonic space for several hours without appearing to substantively aggregate or incorporate into matrix. Eventually, cuticle collagens gradually incorporate and then transiently co-exist in the matrix with pre-cuticle proteins. After the collagens have incorporated, pre-cuticle proteins are removed by endocytosis.

This sequence of events suggests a revision of the classic model for *C. elegans* cuticle assembly, in which sequentially-deposited distinct layers are pushed progressively further away from the plasma membrane [62,63]. The pre-cuticle does not become the outer layer of the mature cuticle. Furthermore, since cuticle collagen matrix assembly does not immediately follow secretion, it could potentially occur external to or within the initial pre-cuticle layer rather than more membrane-proximally. This means that the earliest expressed collagens need not necessarily join the matrix first nor ultimately define more external cuticle layers. For example, despite the early SQT-3 secretion shown here, *sqt-3* mutants are defective in formation of the basal striated layer of the L1 cuticle, which forms only after embryo elongation [55]. We propose that the layered organization of the final cuticle structure is determined not only by the initial timing of cuticle collagen expression and extracellular release, but also by processing events and protein-protein interactions that occur in the extracellular environment or along the still poorly understood routes that these collagens take through the secretory pathway.

We do not currently know if the early pool of secreted collagen has a function or if the collagen that joins the cuticle comes from that extracellular pool (an “outside in” assembly direction) or from a later wave of secreted protein (an “inside out” assembly direction). A very interesting recent study suggested that mammalian fibrillar collagens are initially secreted in soluble form and then re-endocytosed and recycled through a distinct secretory pathway before being competent for fibril elongation [64]. Our observations could be consistent with such a model and set the stage for more detailed future studies of collagen trafficking and matrix assembly in the *C. elegans* system.

### Roles for N-terminal and C-terminal processing in *C. elegans* cuticle collagen secretion and matrix assembly

*C. elegans* cuticle collagens differ from mammalian fibrillar collagens (and are instead more similar to transmembrane MACIT collagens) in having predicted N-terminal cleavage sites that match the consensus for furin/PCSKs rather than ADAMTS proteinases [13,14]. A recent bioinformatic analysis found that 109 of the 173 predicted cuticle collagens contain an N-terminal CFCS that specifically matches the sequence RxxR [24]. Such sequence predictions suggest that a minority of *C. elegans* cuticle collagens are secreted using a conventional N-terminal signal sequence, and that most are predicted to be secreted in type II orientation with a cytosolic N-terminus followed by a transmembrane domain. An RxxR sequence was found in most cuticle collagens with predicted secretion signal sequences (45/56) and in more than half of the predicted transmembrane cuticle collagens (64/117). N-terminal cleavage at CFCS sites is important for function of multiple collagens, since mutations in these sites cause cuticle abnormalities [13,14,22](this work). BLI-4 previously was speculated to be the PCSK that performs these N-terminal cleavages [22,44] and our data strongly support that model for the two collagens tested here, SQT-3 and DPY-17.

Surprisingly, loss of BLI-4 or mutation of its predicted collagen target sites caused a substantial portion of SQT-3 and DPY-17 to aggregate within an apical compartment rather than being released in soluble form to the external environment. This defect cannot be attributed simply to failure to release a transmembrane form of SQT-3 collagen, since a CFCS mutation in DPY-17 (a secreted collagen) also caused aggregation. Furthermore, these aggregates appeared very early, hours before the normal time of cuticle assembly, and they formed independently of the predicted C-terminal proteinase DPY-31. We hypothesize that aggregation occurs as the un-processed procollagens move through the secretory pathway and encounter other partners or environments that allow them to initiate matrix assembly prematurely (Fig. 10E). This model implies that N-terminal processing of SQT-3 and DPY-17 inhibits a key step of matrix assembly during trafficking, in contrast to the more typical scenario where proprotein cleavage facilitates matrix assembly.

In humans, failure to remove the Type I procollagen N-terminus leads to Ehlers-Danos syndrome type VII, a matrix disorder characterized by frequent joint dislocations and tissue fragility [4,9,11]. In that case, the collagen molecules that retain their N-termini incorporate into abnormal fibrils, but to our knowledge no defects in secretion or in the timing of fibril formation have been reported [20,21]. Part of this difference could be technical based on the *in vivo* imaging vs. *in vitro* biochemical approaches used, but it is also likely that there are real biological differences in collagen regulation between different families of collagens. Indeed, in mammalian fibrillar collagens, triple helix formation initiates at the C-terminus and then proceeds towards the N-terminus, whereas in MACIT collagens the opposite is true [15,65]. The direction of trimer assembly is not known for *C. elegans* cuticle collagens, but it has been noted that some have N-terminal coiled-coil regions that could serve as oligomerization domains [66]. *C. elegans* and mammalian collagens also differ in other processing events, for example *C. elegans* cuticle collagens display tyrosine-based crosslinking rather than hydroxylysine based crosslinking [67,68]. It is not unreasonable to hypothesize that N-terminal cleavage may serve different roles in such different contexts.

Our data also suggest that the role of C-terminal cleavage could vary among collagen subtypes. Although SQT-3 contains a predicted BMP1/astacin cleavage site and requires DPY-31 for timely matrix assembly [41,42] (this work), we note that many other cuticle collagens, including DPY-17, do not have recognizable sites for C-terminal processing. Furthermore, C-terminal GFP tags are retained on DPY-17 and multiple other cuticle collagens in matrix structures [60,69], suggesting that those collagens may not undergo such processing. Instead, there may be other unknown modifications, partners, and/or environmental conditions that must be present in order for those collagens to initiate matrix assembly.

### Intracellular vs. extracellular processing of collagens

Despite the clinical importance of Type I procollagen processing, there is still debate in the literature about when and where this happens. C-terminal procollagen cleavage is thought to occur in a late secretory compartment or at the plasma membrane, while N-terminal cleavage may occur either before or after that [3,7]. Growing evidence suggests N-terminal processing occurs at least in part within an ER or Golgi compartment, since the cleaved collagen and isolated N-terminal region can be detected in those locations and since the golgin Giantin is important for collagen processing [18,70–72]. However, uncleaved procollagen and N-proteinase activity also can be readily detected extracellularly in cell culture [4,19] and the best understood N-terminal proteinases, of the ADAMTS family, are secreted proteins also found extracellularly [73,74]. These latter observations led to the traditional view that N-terminal processing occurs outside the cell. It is possible that processing normally occurs in both locations [3] and/or that partially processed and secreted collagens traffic through endocytic recycling compartments before final processing and matrix assembly [64].

Our data are more consistent with intracellular N-terminal cleavage of *C. elegans* cuticle collagens, since BLI-4 is mainly detected intracellularly and its loss leads to intracellular aggregates of SQT-3 and DPY-17. In most cases, the aggregates appear to extend near to and potentially across the apical plasma membrane, suggesting that they could be present in tubulovesicular compartments that have access to the outside environment. Future identification of the BLI-4- and aggregate-containing cellular compartments, combined with direct assays for cleavage and other modifications, should more precisely define the site and order of cuticle collagen processing. Ultimately, these studies should reveal how processing events help control the time and place of collagen assembly to construct the various cuticle structures and layers observed *in vivo*.

## Methods

### Strains and animal husbandry

See Table S1 for a list of all strains used in this work. *C. elegans* N2 was used as the wild-type strain. Unless otherwise indicated, strains were grown at 20°C under standard conditions [75]. See Table S2 for specific mutant lesions and a list of all sgRNAs and primers used for genome editing or transgenics. Tagged collagen and BLI-4::SfGFP strains and CFCS mutants of *sqt-3* and *dpy-17* (RxxR to AxxA) were made by Suny Biotech (Fuzhou, China). The endogenous fusions are functional based on viability and normal body morphology of the homozygotes. Rescue transgene *csEx919 (bli-4+)* was generated by microinjection of fosmid WRM069bE05 (20 ng/ul) with *sur-5::GFP* (30 ng/ul) and bluescript SK+ (50 ng/ul).

### Generation of *bli-4* alleles

Although *bli-4* null alleles had already been described [44,76], they were generated on chromosomes carrying other markers that could complicate analysis; therefore, we opted to generate new alleles in an N2 background. Mutant alleles were generated by CRISPR-Cas9 genome editing using methods described in [77] and the sgRNAs listed in Table S2. N2 hermaphrodites were injected with sgRNAs (IDT), Cas9 (University of California Berkeley), and the marker pRF4, and F2 progeny were screened for expected embryonic or larval lethal phenotypes. Mutant alleles were recovered over a genetic balancer (either *hT2 (I;III)* or *szT1 (I;X)*) and then rescued with *bli-4+* transgene *csEx919*. Mutant lesions were identified by PCR amplification and Sanger sequencing. We originally identified 4 putative *bli-4* null alleles, 8 putative *bli-4(c/d)* alleles, and 3 putative *bli-4(d)* alleles; however, only the alleles described here had small (<50 bp) deletions that permitted PCR-amplification with our methods, while the remaining alleles appeared to have larger deletions or rearrangements and were not further characterized.

### Microscopy and image processing

For timelapse imaging, ~24 cell stage embryos were mounted in egg buffer/methyl cellulose with 20μM beads as spacers [78], incubated at 20° C for 4 hours, and then imaged using a stage temperature controller at 12° C and a Leica TCS SP5 confocal (20 z-planes at 0.5 μM spacing and 15 minute time spacing, total 16 hours). To immobilize animals for still imaging, embryos or larvae were suspended in M9 buffer with 10mM levamisole and mounted on 2% agarose pads supplemented with 20mM sodium azide. DIC and epifluorescence images were obtained with a Zeiss Axioskop (Carl Zeiss Microscopy) fitted with a Leica DFC360 FX camera with Qcapture (Qimaging) software. Confocal images were captured with a Leica TCS SP8 confocal microscope, except for images in Figs. 9C,D and 10C,D, which were captured with a Zeiss LSM780 confocal microscope. Images were analyzed and processed in FIJI [79]. To quantify fusion protein accumulation in the extraembryonic space, fluorescence intensity was measured in a 2×2 μm region within a single confocal slice of each specimen. For permeability assays, L1 larvae were incubated in 2 μg/ml Hoescht dye 33258 (Sigma) in M9 buffer for 15 minutes at room temperature, then washed twice with M9 before imaging.

### Statistical Analyses

Statistical analyses were performed using GraphPad Prism. In all dot-plots, lines and error bars indicate the mean and standard deviation, respectively, and each dot represents a measurement from a single animal. To perform statistical analyses on quantitative measurement data, genotypes were compared using a non-parametric Mann-Whitney test. To perform statistical analyses on categorical data, phenotypes were classified as either normal or abnormal and then proportions compared using a two-tailed Fisher’s Exact test.

### *bli-4* isoform analysis from single-cell RNA-seq data

scRNA-seq reads from [49] were remapped to each exon of the protein coding genes in Wormbase genome build WS277 using cellranger software (10x genomics). The resulting UMI counts per exon for each tissue were combined in VisCello software with the existing *C. elegans* embryo scRNA-seq atlas [49]. Pseudobulk expression levels of each *bli-4* exon in reads per million were calculated in cells annotated as each cell type, and in major tissue classes as annotated in [49]. Code used for these analyses are available at Github: https://github.com/jisaacmurray/bli4_paper.

### Western blots

Western blots with BLI-4::SfGFP(int) were performed as in [80] using primary antibody goat anti-GFP (Rockland Immunochemicals, 600-101-215, 1:1000 dilution) and secondary antibody anti-goat-HRP (Rockland Immunochemicals, 605-4302, 1:5000 dilution). Embryos were collected from gravid adults via the alkaline bleach method and then allowed to develop for 5 hours in M9 before processing.

## Acknowledgements

We thank Alison Frand and Michel Labouesse for strains, Andrea Stout and the Microscopy core at University of Pennsylvania for training and assistance with imaging, and Priya Sivaramakrishnan for assistance with time-lapse imaging. We thank Nathalie Pujol, Helen Schmidt, Nicholas Serra, and Trevor Barker for helpful discussions and comments on the manuscript, and Nathalie Pujol for hosting M.V.S. during the final stages of its preparation. Some strains used in this study were provided by the *Caenorhabditis* Genetics Center, which is funded by the NIH Office of Research Infrastructure Programs (P40 OD010440). This work was supported by National Institutes of Health grants R35GM136315 to MVS, R35GM134970 to ADC, and training award T32 AR007465 to JDC.

## Supplemental Materials

**Table S1. *C. elegans* strains used**

**Table S2. Sequences of *bli-4* mutant alleles and insertions**

**Figure S1. Pre-cuticle mCherry fusions accumulate in lysosome-like compartments following endocytosis**

A,B) Pre-cuticle mCherry fusions, but not SfGFP fusions, mark large internal structures in late embryos (4 hours after 1.5-fold). A) *noah-1(mc68* [NOAH-1::mCherry(int)]) compared to *aaaIs25* [NOAH-1::SfGFP(int)]. B) *lpr-3(cs266* [ss::mCherry::LPR-3]) compared to *lpr-3(cs250* [ss::SfGFP::LPR-3]). All images are maximum intensity projections from confocal Z-slices and representative of at least 5 embryos per genotype. Scale bar, 10 microns.

**Figure S2. *bli-4* isoform expression in the embryo**

Embryo scRNA-seq reads from [49] were mapped to the 3’ ends of the various *bli-4* isoforms to estimate expression levels in the major tissue classes (Methods). RPM, reads per million. Our methods could not distinguish isoforms f and g, despite their different 3’ coding regions, since these isoforms share a similar 3’ UTR. The most highly expressed isoform in the embryo and in the epidermis was the CRD-containing isoform, *bli-4d*, with more modest levels of *bli-4a* and f*/g,* and negligible evidence for other isoforms.

**Figure S3. BLI-4 is dispensable for trafficking and apical membrane localization of the ZP protein NOAH-1**

A) Schematic diagram of the NOAH-1::SfGFP(int) transgene fusion (*aaaIs25*) [82], which was used instead of the endogenous fusion because of chromosomal linkage of the *bli-4* and *noah-1* loci. The SfGFP tag (green triangle) is located between the Plasminogen (PAN) and ZP domains, as indicated. The C-terminal CFCS (RKKR, orange) is located before a predicted transmembrane (TM) domain. B) NOAH-1::SfGFP(int) appears similar between *WT* and *bli-4* mutant embryos (1.5-fold + 3 hour stage). Images are single confocal Z-slices and are representative of at least 10 embryos per genotype. C) Peak membrane fluorescence intensity was calculated with FIJI [79] using line scans across single confocal Z-slices, as shown in B. There was no significant difference (ns) among the genotypes.

**Supplementary videos 1 and 2**

Time-lapse videos of an elongating *bli-4(cs281)* embryo and rescued sibling, taken at 12°C. Videos are representative of at least 5 imaged specimens per genotype.

