## Supplementary figures and images for "The proprotein convertase BLI-4 promotes collagen secretion during assembly of the *Caenorhabditis elegans* cuticle"

### Supplemental Figure 1

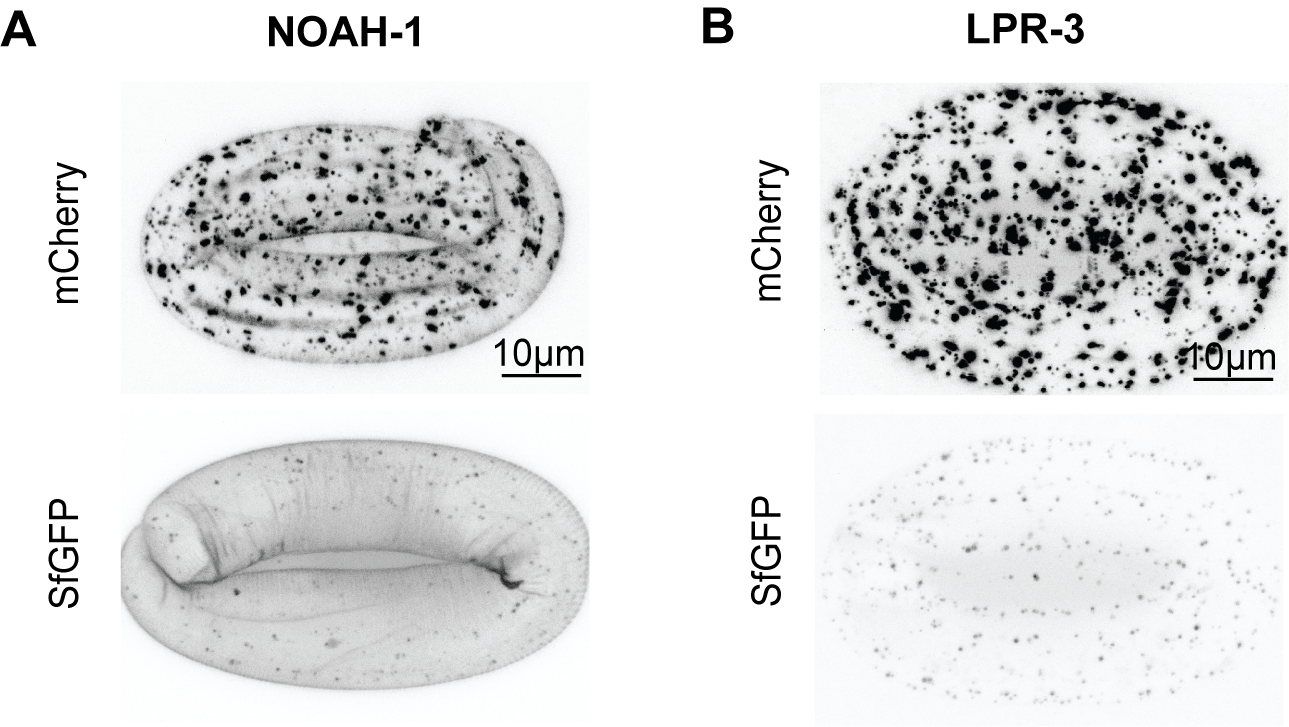

### Supplemental Figure 2

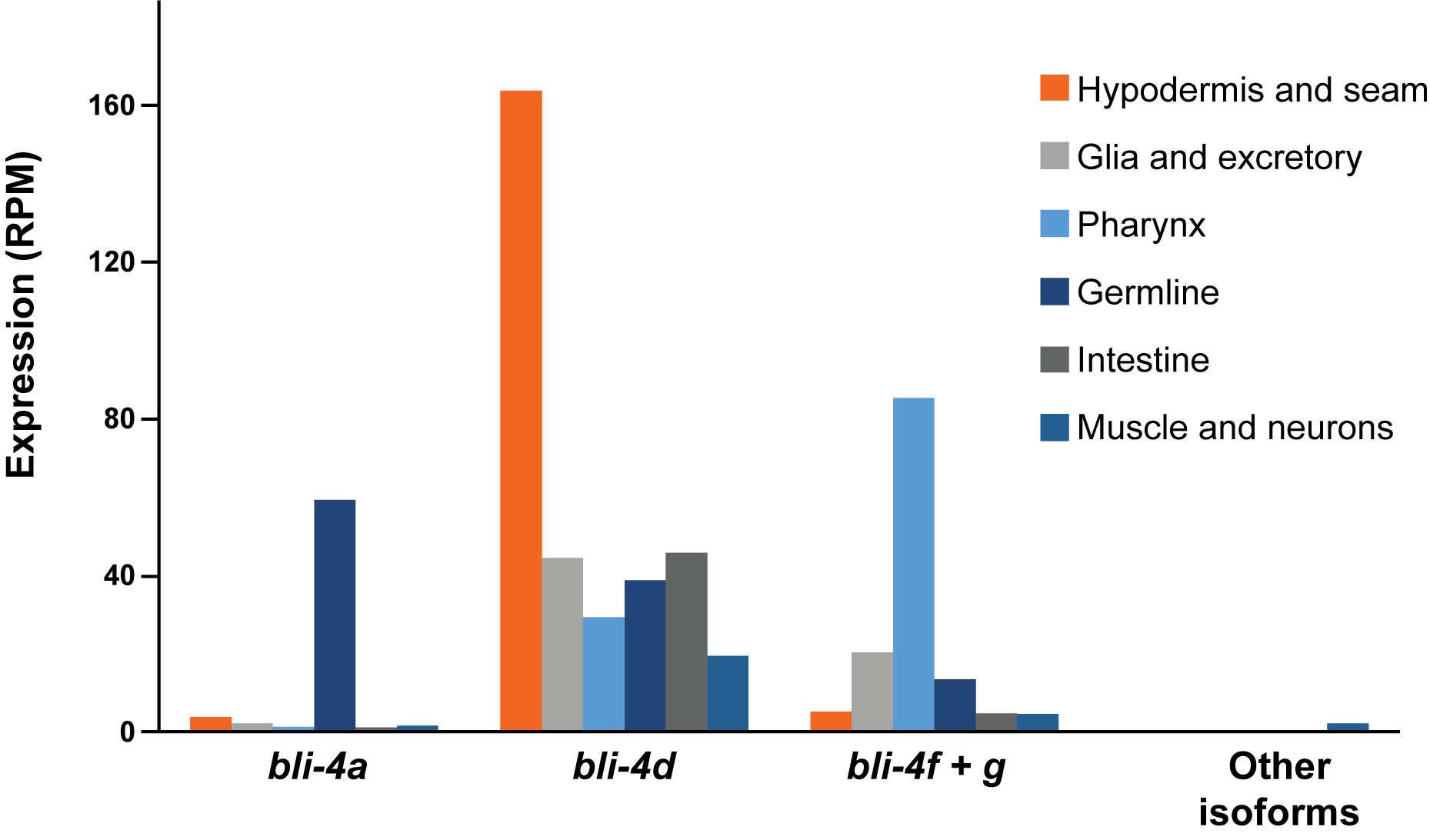

### Supplemental Figure 3

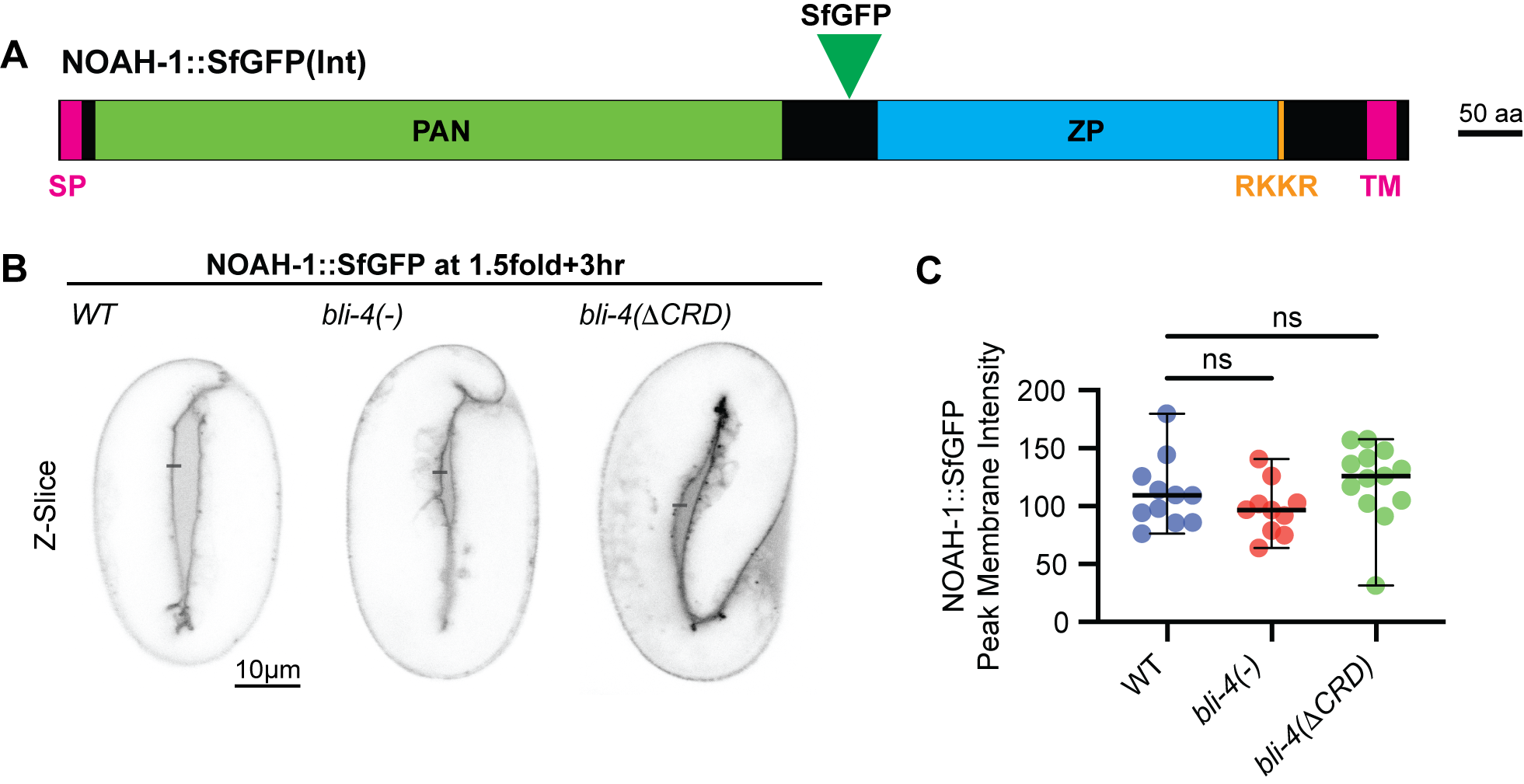
